# A mosaic-type trimeric RBD-based COVID-19 vaccine candidate induces potent neutralization against Omicron and other SARS-CoV-2 variants

**DOI:** 10.1101/2022.03.29.486173

**Authors:** Jing Zhang, Zi Bo Han, Yu Liang, Xue Feng Zhang, Yu Qin Jin, Li Fang Du, Shuai Shao, Hui Wang, Jun Wei Hou, Ke Xu, Ze Hua Lei, Zhao Ming Liu, Jin Zhang, Ya Nan Hou, Ning Liu, Fu Jie Shen, Jin Juan Wu, Xiang Zheng, Xin Yu Li, Xin Li, Wei Jin Huang, Gui Zhen Wu, Ji Guo Su, Qi Ming Li

## Abstract

Large-scale populations in the world have been vaccinated with COVID-19 vaccines, however, breakthrough infections of SARS-CoV-2 are still growing rapidly due to the emergence of immune-evasive variants, especially Omicron. It is urgent to develop effective broad-spectrum vaccines to better control the pandemic of these variants. Here, we present a mosaic-type trimeric form of spike receptor-binding domain (mos-tri-RBD) as a broad-spectrum vaccine candidate, which carries the key mutations from Omicron and other circulating variants. Tests in rats showed that the designed mos-tri-RBD, whether used alone or as a booster shot, elicited potent cross-neutralizing antibodies against not only Omicron but also other immune-evasive variants. Neutralizing antibody titers induced by mos-tri-RBD were substantially higher than those elicited by homo-tri-RBD (containing homologous RBDs from prototype strain) or the inactivated vaccine BBIBP-CorV. Our study indicates that mos-tri-RBD is highly immunogenic, which may serve as a broad-spectrum vaccine candidate in combating SARS-CoV-2 variants including Omicron.

## Introduction

The severe acute respiratory syndrome coronavirus 2 (SARS-CoV-2) is continuously evolving, and the emergence of new variants has caused successive waves of coronavirus disease 2019 (COVID-19). Among the circulating variants, five strains, including Alpha, Beta, Gamma, Delta, and Omicron, have been classified into variants of concern (VOCs) by the world health organization (WHO) (https://www.who.int/en/activities/tracking-SARS-CoV-2-variants/). Alpha, as the first VOC, became a globally dominant strain in early 2021, which was then replaced by Delta variant from the summer of 2021. These two variants exhibited slightly and moderately less sensitivity to neutralization by serum from vaccinated individuals, with 1-2-fold and 2-3-fold reductions in neutralizing titers, respectively (Lipsitch et al., 2022; Muik et al., 2021; Widge et al., 2021; Planas et al., 2021; Wu et al., 2021b; Pegu et al., 2021). Beta and Gamma variants outbroke in Africa and South America from early to mid-2021, respectively. Beta variant showed significantly greater immune escape capability and was 3-15-fold less susceptible to neutralization by vaccine-induced antibodies (Lipsitch et al., 2022; Planas et al., 2021; Edara et al., 2021; Shen et al., 2021). Gamma strain also exhibited obvious reduced neutralizing sensitivity, but the reduction in neutralizing titers was not as substantial as that of Beta strain (Lazarevic et al., 2021; Wang et al., 2021; Dejnirattisai et al., 2021). Besides that, the variants of interest (VOIs) designated by WHO, including Lambda and Mu, have also been reported to have certain immune evasion abilities (Liu et al., 2021; Lou et al., 2021). The potential immune escape of SARS-CoV-2 variants raised concerns about the efficacy of current COVID-19 vaccines, and new generation vaccines specific to Beta variant have been developed by several groups (Wu et al., 2021a; Callaway and Ledford, 2021; Logue et al., 2021).

Recently, the pandemic of Omicron variant posed a more serious threat to the protective effectiveness of the currently used vaccines. The Omicron variant, also known as B.1.1.529, was first detected in Botswana and reported from South Africa, which was considered to be associated with the sharp rise of infection cases in multiple provinces in South Africa (https://www.who.int/news/item/26-11-2021-classification-of-omicron-(b.1.1.529)-sars-cov-2-variant-of-concern). Preliminary evidence indicates that this variant may be more transmissible and may have a higher reinfection risk than other VOCs, and thus just two days after its discovery, Omicron has been assigned as a VOC by the WHO (Vaughan, 2021; Callaway, 2021; Kupferschmidt and Vogel, 2021; Torjesen, 2021). According to the data from GISGAID, so far, Omicron has spread rapidly to more than 95 countries and the reported cases of this variant are growing rapidly around the world (https://www.gisaid.org/hcov19-variants/). Omicron carries 15 mutations in the receptor binding domain (RBD) of the spike (S) protein that is the immunodominant target of neutralizing antibodies. Compared with other VOCs, the substantially more mutations, as well as their important locations for antibody binding, may enable Omicron to escape the immune protection offered by previous infection or vaccination. Our recent study on the neutralizing sensitivity of the convalescent serum against pseudo-typed Omicron showed that the neutralizing titer was significantly decreased 8.4-fold compared to the D614G strain (Zhang et al., 2022). Several studies reported by other groups also demonstrated remarkable resistance of Omicron to neutralization by sera from convalescent patients or vaccinated individuals. Neutralizing antibody titers against Omicron variant in the individuals administered by mRNA COVID-19 vaccines were dramatically reduced 8.6-22 folds compared to the D614G reference strain and by contrast, only 4.3-5-fold decline was observed for Beta variant (Liu et al., 2022; Cele et al., 2022). Sera from the individuals vaccinated with two doses of ChAdOx1 or Ad26.COV2.S failed to neutralize Omicron (Liu et al., 2022; Rössler et al., 2021). Tests on a panel of existing SARS-CoV-2 neutralizing antibodies showed that Omicron variant evaded neutralization of the majority of these antibodies (Cao et al., 2022). As of February 2022, the Omicorn variant has evolved into three lineages: BA.1, BA.2 and BA.3, and recently, the mutations from Omicron and Delta variants were combined to create a Deltacron super variant. The extensive immune-escape capability of Omicron and other circulating SARS-CoV-2 variants from previous infections and vaccinations raises an urgent need of developing effective broad-spectrum vaccines against these immune-evasive variants.

Guided by structural and computational analyses of S protein RBD, we have developed a trimeric form of RBD vaccine candidate, i.e., the homologous trimeric RBD (homo-tri-RBD), in which three RBDs from the prototype SARS-CoV-2 strain were connected end-to-end into a single molecule and co-assembled into a trimeric form (Liang et al., 2022). Animal experiments and clinical trials have demonstrated potent protections offered by this vaccine against SARS-CoV-2. At present, homo-tri-RBD has completed phase I/II clinical trial and approved by the United Arab Emirates for emergency use. Here, our vaccine design scheme was extended to broaden its immune response against SARS-CoV-2 variants. Targeting SARS-CoV-2 Omicron and other immune-evasive variants, we present a mosaic-type trimeric RBD (mos-tri-RBD) vaccine candidate, in which the key mutations derived from Omicron (BA.1) as well as other VOCs and VOIs were integrated into the immunogen. The immunogenicity of the designed mos-tri-RBD was evaluated in rats by using live-virus neutralization assays. To illustrate its superiority in stimulating broad-spectrum neutralizing activities against Omicron and other immune-evasive variants, the cross-reactive immunity induced by mos-tri-RBD was compared with that elicited by homo-tri-RBD and the inactivated vaccine BBIBP-CorV. Especially, given that large-scale populations worldwide have received the primary series of vaccination, the immunogenicity of the designed mos-tri-RBD as a booster dose following the primary vaccination of BBIBP-CorV was also evaluated and compared with the booster vaccinations of homo-tri-RBD and BBIBP-CorV. Our results showed that the immunization with mos-tri-RBD either alone or as a booster dose elicited potent broad-reactive neutralizing response against SARS-CoV-2 variants including Omicron, which was immunogenically superior to homo-tri-RBD and BBIBP-CorV.

## Results

### Design of the mos-tri-RBD based on the RBDs from SARS-CoV-2 Omicron and other variants

RBD forms a relatively compacted and isolated domain in the structure of spike (S) protein, and the beta-sheets in the core as well as the existence of four disulfide bonds stabilizes the tertiary structure of the domain. The N- and C-termini of RBD are close together, and there exist long loops in both termini. These structural features inspired our construction of a trimeric form of RBD (tri-RBD) through an end-to-end connection of three RBDs into a single chain, in which their own long loops at the N- and C-termini serve as the linkers. The designed tri-RBD enables the accommodation of three RBDs in one immunogen, which can be extended to include Omicron RBD into the immunogen.

In this study, mos-tri-RBD was designed targeting Omicron as well as other emerging variants with distinct immune evasion capability. Mos-tri-RBD also consisted of three RBDs, one of which was derived from Omicron and the other two were artificially designed to carry the key mutations appearing in SARS-CoV-2 variants. These key mutations integrated into mos-tri-RBD were selected as those appearing in SARS-CoV-2 VOCs or VOIs and simultaneously being ranked in the top ten most frequently occurring mutations in RBD as counted by Wei group (https://weilab.math.msu.edu/MutationAnalyzer/). According to this criterion, a total of eight mutations were chosen and introduced into the two artificially designed RBDs (The details on these mutations were provided in Table 1), where one RBD contained the mutations of K417N, L452R, T478K, F490S and N501Y, and the other included K417T, S477N and E484K (Figure 1A). Many pieces of evidence have indicated that these mutations largely contribute to the immune escape of the related variants. We sought to integrate these key mutations into a single immunogen to elicit cross-neutralizations against not only SARS-CoV-2 Omicron but also other circulating variants. To facilitate the self-trimerization of mos-tri-RBD, for each RBD the residues 319-537 were truncated from the S protein to retain the long loops at both termini, as shown in Figure 1A. Our previous studies have shown that the RBDs with this truncation scheme can correctly fold and co-assemble into a trimeric structure (Liang et al., 2022).

**Table 1.** The details on the eight selected mutations integrated into the two artificially designed RBDs.

| Mutations | Rank of the observed frequency | VOCs and VOIs carrying the mutations |
| --- | --- | --- |
| L452R | 1 | Delta |
| T478K | 2 | Delta、Omicron |
| N501Y | 3 | Alpha、Beta、Gamma、Omicron、Mu |
| E484K | 4 | Beta、Gamma、Mu |
| K417T | 5 | Gamma |
| S477N | 6 | Omicron |
| K417N | 8 | Beta、Omicron |
| F490S | 10 | Lambda |

**Figure 1.**
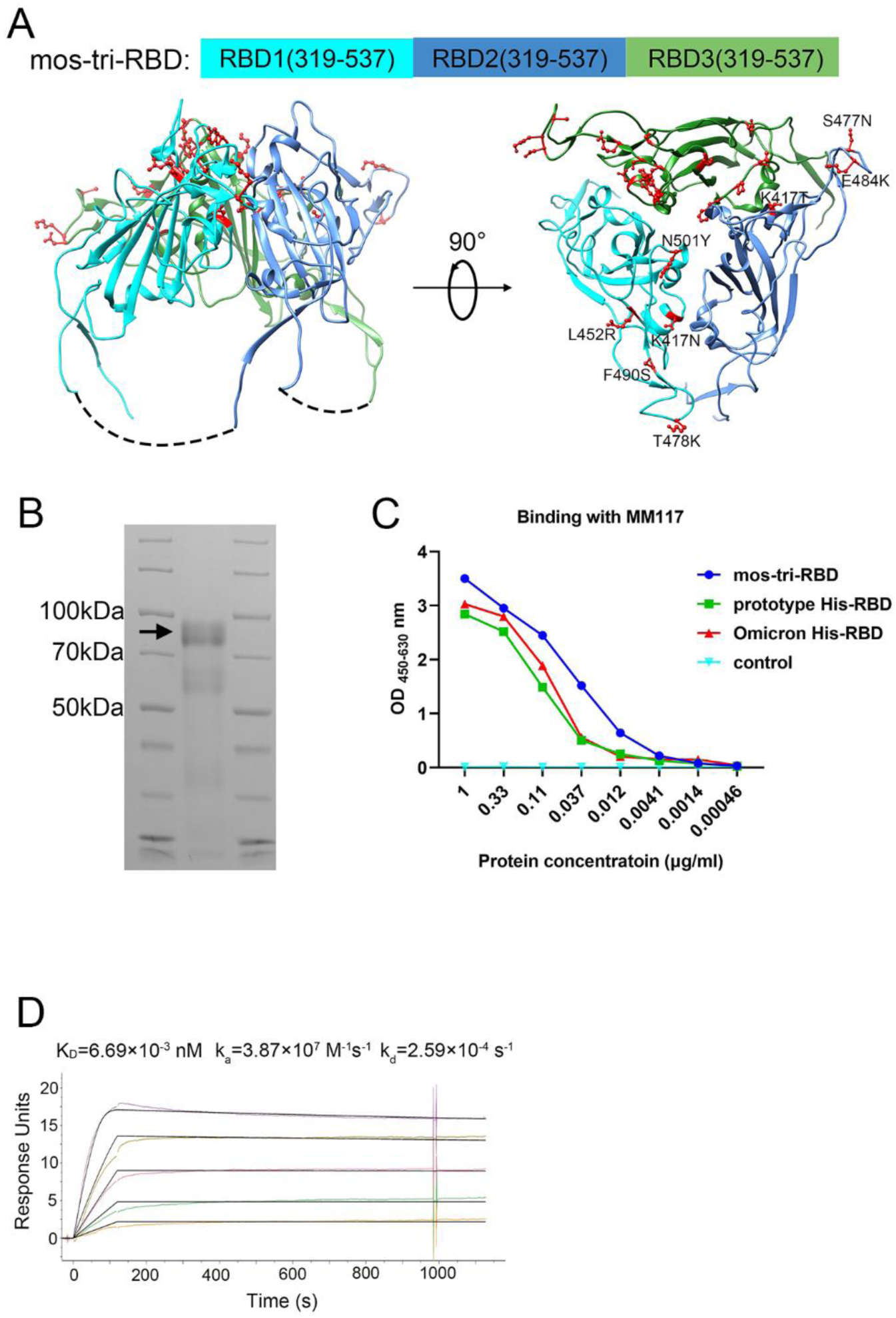
Design, expression and characterization of the mosaic-type trimeric form of RBD (mos-tri-RBD). **(A)** Schematic illustration of the designed mos-tri-RBD. In mos-tri-RBD, three heterologous RBDs were connected end to end into a single chain and co-assembled into a trimeric structure. For the three RBDs, one was derived from the Omicron (BA.1) variant (green color), and the other two were artificially designed harboring the key immune-evasion-related mutations that emerged in SARS-CoV-2 variants, in which one contained the mutations of K417N, L452R, T478K, F490S and N501Y (cyan color), and the other one contained K417T, S477N and E484K (blue color). These mutations are highlighted in the red ball-and-stick model in the figure. Each RBD subunit in mos-tri-RBD was composed of the residues 319-537 from the spike protein. The dotted curves in the figure represent the direct connection between the C-terminus of the former RBD and the N-terminus of the latter RBD. The schematic structure of mos-tri-RBD was drawn by Chimera software (Pettersen et al., 2004) based on the PDB file with accession number 6zgi. **(B)** SDS-PAGE analysis of the recombinant mos-tri-RBD. **(C)** Concentration-dependent binding ability of mos-tri-RBD with an RBD-specific monoclonal neutralizing antibody MM117 tested using ELISA. **(D)** Binding avidity of mos-tri-RBD with the receptor hACE2 measured using SPR assay. **Figure 1 – source data 1** **The raw files of SDS-PAGE results**. **Figure 1 – source data 2** **Concentration-dependent binding ability of mos-tri-RBD with the antibody MM117**.

### Expression, identification and characterization of the recombinant mos-tri-RBD protein

The recombinant mos-tri-RBD was expressed using CHO cells, and then purified by chromatography and ultrafiltration as described previously (Liang et al., 2022). SDS-PAGE analysis displayed an obvious band corresponding to the molecular mass of about 92 kDa (Fig. 1B), implying the formation of the trimeric RBD. The measured mass was larger than the theoretical value calculated by the sequence, which was attributed to the heavy glycosylation of the protein as discussed in our previous studies (Liang et al., 2022). To determine the biological function of the recombinant mos-tri-RBD, its binding ability to an RBD-specific monoclonal neutralizing antibody MM117 was tested using enzyme-linked immunosorbent assay (ELISA). MM117 has been proved to be able to bind specifically with the RBDs of SARS-CoV-2 prototype, Omicron and Delta strains. Protein concentration-dependent binding activity was observed for MM117, suggesting the formation of native conformation of the RBDs in mos-tri-RBD (Fig. 1C). Furthermore, the binding avidity of mos-tri-RBD with the receptor human angiotensin converting enzyme 2 (hACE2) was also measured using surface plasmon resonance (SPR) assay. The association rate constant (*k*_a_) and dissociation rate constant (*k*_d_) were quantified to be 3.87 × 10^7^ *M*^−1^*s*^−1^ and 2.59 × 10^−4^ *s*^−1^, respectively, and thus the apparent dissociation constant *K*_D_ was determined to be 6.69 × 10^−3^ *nM* (Fig. 1D). The results of SPR assay implied that the recombinant mos-tri-RBD can recognize hACE2 specifically with high avidity, which verified the functionality of the designed protein and the correct folding of each RBD into its native conformation. All these results suggested that the designed mos-tri-RBD assembled into a trimeric form and each RBD subunit correctly folded into its native structure.

### Mos-tri-RBD induced potent cross-reactive neutralizing response against the live viruses of SARS-CoV-2 Omicron and other immune-evasive variants

To evaluate the cross-reactive immunogenicity of the designed mos-tri-RBD, we intramuscularly immunized rats using two doses of mos-tri-RBD mixed with Aluminum adjuvant three weeks apart. Another three groups of rats received two doses of homo-tri-RBD, BBIBP-CorV or adjuvant, respectively, with the same immunization regimen were used for comparison. Sera from the immunized rats were collected on day 7 after the last vaccination (Fig. 2A). Neutralizing antibody titers in the sera against multiple SARS-CoV-2 strains, including prototype, Omicron, Beta and Delta strains, were detected using live-virus neutralization assay.

**Figure 2.**
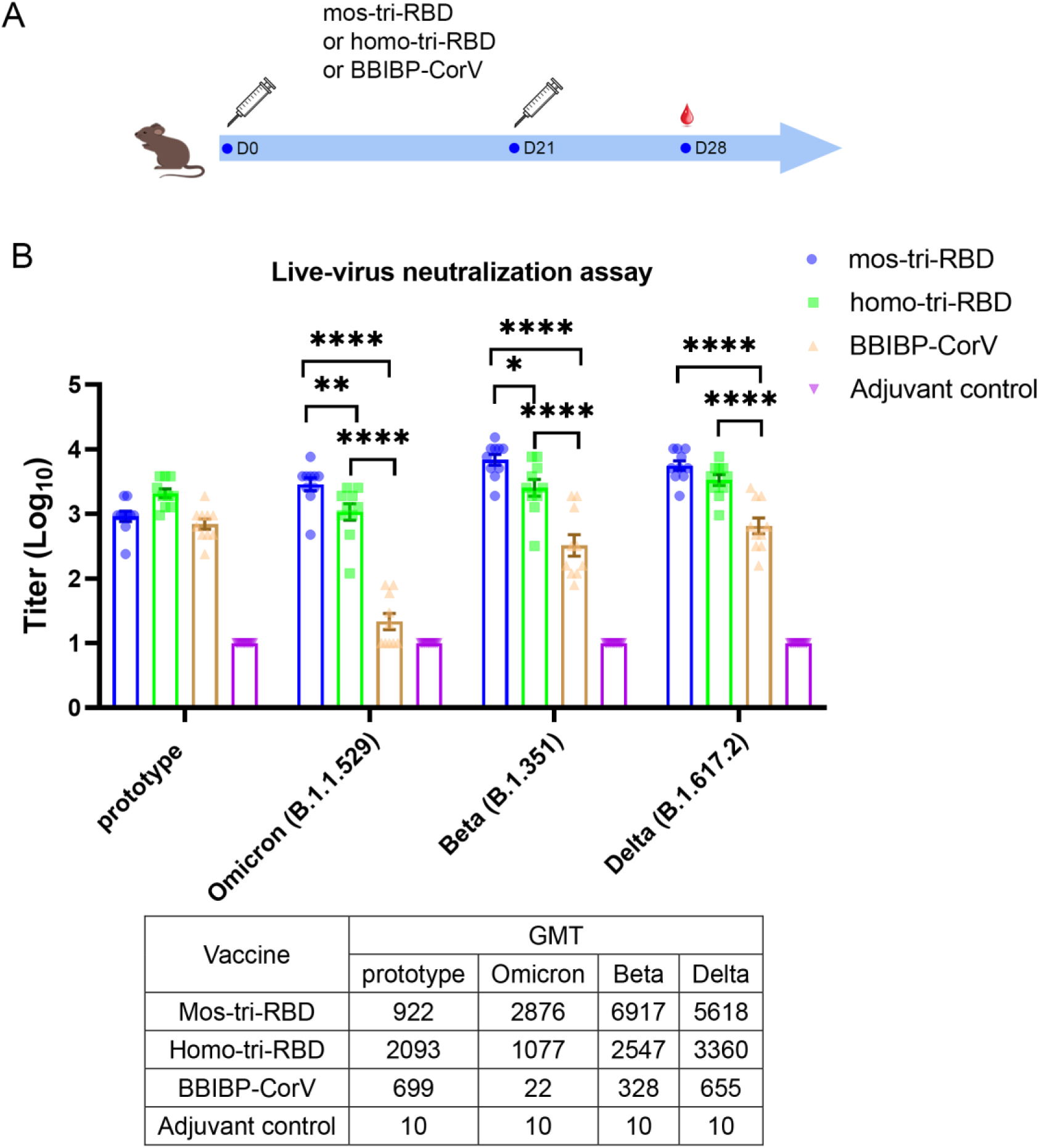
Evaluation of the cross-reactive immunogenicity of mos-tri-RBD against multiple SARS-CoV-2 strains, including prototype, Omicron, Beta and Delta strains, using live-virus neutralization assay. **(A)** Timeline of rat immunization and serum collections. Rats were immunized intramuscularly with two doses of mos-tri-RBD with three weeks apart. Another three groups of rats received two doses of homo-tri-RBD, BBIBP-CorV and adjuvant, respectively, were used for comparison. Sera from all the immunized rats were collected on day 7 after the last vaccination. **(B)** Neutralizing antibody titers elicited by mos-tri-RBD compared with those elicited by homo-tri-RBD and BBIBP-CorV against the live-viruses of SARS-CoV-2 prototype strain, and Omicron, Beta and Delta variants. Data are presented as mean ± SEM. One-way ANOVA followed by the LSD t-test was used for the comparison of data between different groups. *P<0.05, **P<0.01, ****P<0.0001. GMT values are displayed in the lower part of the figure. **Figure 2 – source data 1** **Individual data of live-virus neutralizing antibody titers against several SARS-CoV-2 circulating strains elicited by mos-tri-RBD compared with those elicited by homo-tri-RBD and BBIBP-CorV**.

Live-virus neutralization assay showed that compared with the prototype virus, Omicron variant exhibit substantially less susceptible to neutralization elicited by two doses of BBIBP-CorV vaccination. The geometric mean titer (GMT) of neutralizing antibodies against prototype strain was 699, whereas the Omicron-specific neutralizing antibody titer in half of the rats was less than the detectable limit of the assay (Figure 2B), suggesting substantial evasion of Omicron variant from the immunity elicited by BBIBP-CorV. The result was consistent with the findings of other studies that Omicron exhibited significant immune escape capability (Zhang et al., 2022; Liu et al., 2022; Cele et al., 2022; Rössler et al., 2021; Cao et al., 2022). Compared with BBIBP-CorV, homo-tri-RBD vaccination significantly improved the neutralizing antibody GMT against Omicron virus from 22 to 1077, with a 49.0-fold increase. Furthermore, remarkably enhanced neutralizing antibody titers against Omicron were elicited by mos-tri-RBD vaccination, in which the neutralizing GMT reached 2876, with 2.7-fold and 130.7-fold increases in comparison to the homo-tri-RBD and BBIBP-CorV vaccinations, respectively (Figure 2B). Statistical analysis showed that the anti-Omicron neutralizing antibody response elicited by mos-tri-RBD was significantly higher than that of homo-tri-RBD (*P*=0.0057) and the neutralization elicited by homo-tri-RBD was also significantly higher than that of BBIBP-CorV (*P*<0.0001). Our study demonstrated that due to containing the RBD from Omicron variant, the designed mos-tri-RBD exhibited much higher immunogenicity against this variant than homo-tri-RBD and BBIBP-CorV. Mos-tri-RBD may serve as an effective vaccine candidate in fighting against Omicron variant.

Similar results were also observed for Beta and Delta variants. In the rats immunized with BBIBP-CorV, the neutralizing antibody GMT against Beta variant was reduced by 2.1-fold in comparison with that against the prototype virus, suggesting a considerable immune escape of the variant. Many previously reported results also found that Beta variant distinctly evaded the immunity offered by natural infection or vaccination (Lipsitch et al., 2022; Planas et al., 2021; Edara et al., 2021; Shen et al., 2021). While, compared to BBIBP-CorV, the anti-Beta neutralizing antibody GMT elicited by homo-tri-RBD was increased from 328 to 2547 (7.8-fold), and further improved to 6917 (21.1-fold) by mos-tri-RBD vaccination (Figure 2B). Similarly, for Delta variant, the neutralizing antibody GMTs induced by homo-tri-RBD and mos-tri-RBD were 5.1-fold and 8.6-fold, respectively, higher than that induced by BBIBP-CorV (Figure 2B). Our results indicated that besides Omicron variant, mos-tri-RBD was also highly immunogenic against Beta and Delta variants.

In summary, live-virus neutralization assays demonstrated that the designed mos-tri-RBD, which integrated key residues from Omicron and other circulating SARS-CoV-2 variants into a single antigen, could serve as a broad-spectrum COVID-19 vaccine candidate against not only Omicron variant but also other SARS-CoV-2 variants. However, it should be noted that due to the absence of wild-type RBD in the mosaic antigen, the neutralizing antibody response against SARS-CoV-2 prototype strain stimulated by mos-tri-RBD was lower than that by homo-tri-RBD, but still comparable to that by BBIBP-CorV (Figure 2B).

### Mos-tri-RBD as a booster dose induced cross-neutralization against the pseudo-typed SARS-CoV-2 Omicron as well as other VOCs and VOIs

Given that large-scale populations worldwide have received the primary series of vaccination, the immunogenicity of the designed mos-tri-RBD as a booster dose was mainly evaluated in this study. Rats were primed with a dose of BBIBP-CorV, and successively boosted with a dose of mos-tri-RBD (“BBIBP-CorV+mos-tri-RBD” group), homo-tri-RBD (“BBIBP-CorV+homo-tri-RBD” group) or BBIBP-CorV (“BBIBP-CorV+BBIBP-CorV” group), as shown in Fig. 3A. Another group of rats received two doses of adjuvant was served as a control. On day 7 post-boost, the sera from the immunized rats were collected, and the neutralizing response against various SARS-CoV-2 strains was tested using pseudo-virus neutralization assays.

**Figure 3.**
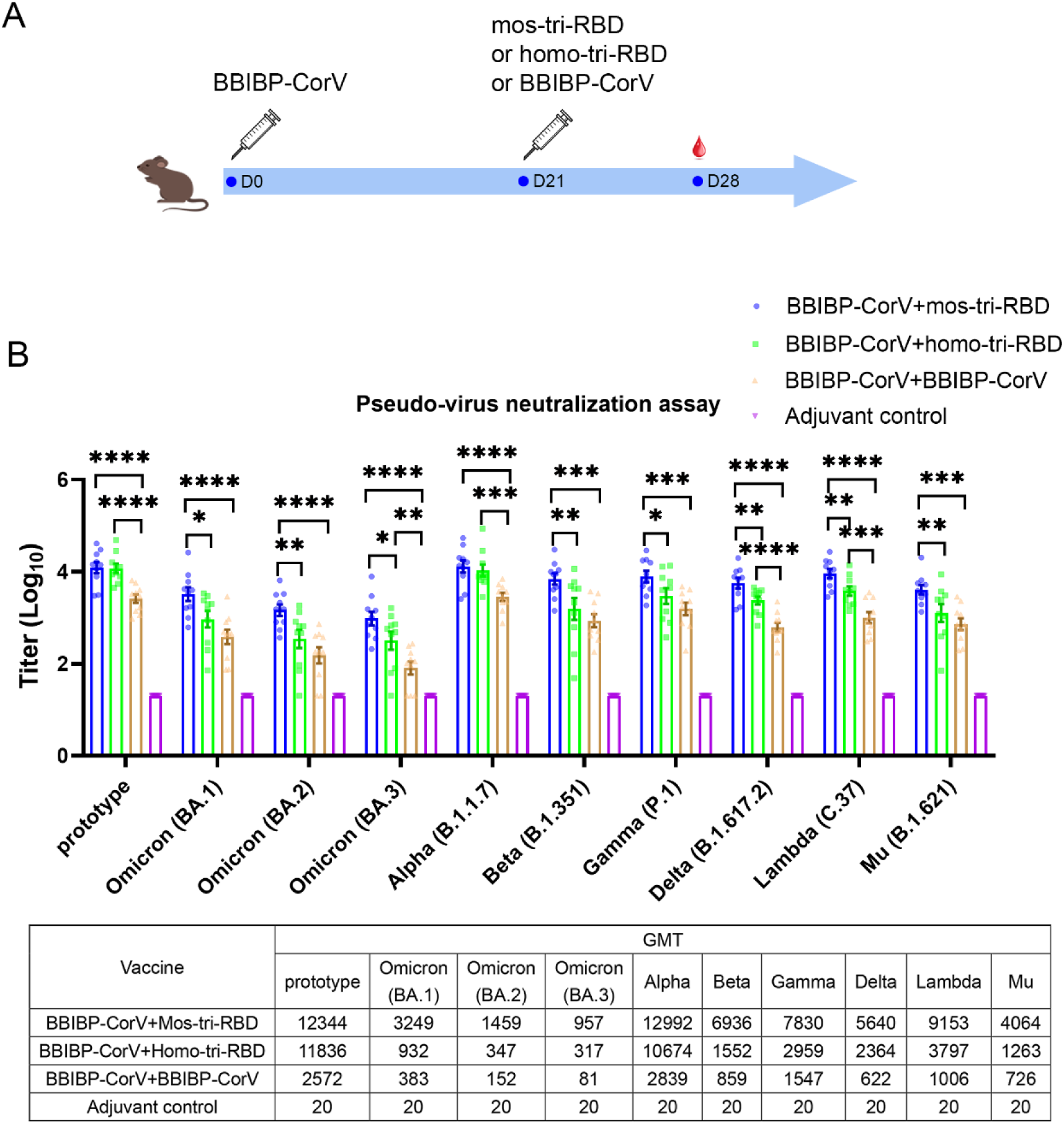
Evaluation of the cross-reactive immunogenicity of mos-tri-RBD as a booster shot against SARS-CoV-2 Omicron as well as other VOCs and VOIs using pseudo-virus neutralization assays. **(A)** Timeline of rat immunization and serum collections. Rats were primed with a dose of BBIBP-CorV and boosted by mos-tri-RBD, homo-tri-RBD or BBIBP-CorV with three weeks apart. Another group of rats vaccinated with two doses of adjuvant served as control. The sera of all the immunized rats were collected on day 7 post-boosting immunization. **(B)** Neutralizing antibody titers elicited by “BBIBP-CorV+mos-tri-RBD” vaccination compared with those elicited by “BBIBP-CorV+homo-tri-RBD” and “BBIBP-CorV+BBIBP-Corv” vaccinations against the pseudo-viruses of SARS-CoV-2 Omicron as well as other VOCs and VOIs. Data are presented as mean ± SEM. One-way ANOVA followed by the LSD t-test was used for the comparison of data between different groups. *P<0.05, **P<0.01, ****P<0.0001. GMT values are displayed in the lower part of the figure. **Figure 3 – source data 1** **Individual data of pseudo-virus neutralizing antibody titers against various SARS-CoV-2 circulating strains elicited by “BBIBP-CorV+mos-tri-RBD” vaccination compared with those elicited by “BBIBP-CorV+homo-tri-RBD” and “BBIBP-CorV+BBIBP-CorV” vaccinations**.

Pseudo-virus neutralization assay demonstrated that all the three prime-boosting vaccinations induced elevated neutralizing antibodies in comparison to the adjuvant control group against the pseudo-virus of SARS-CoV-2 prototype strain. Although mos-tri-RBD did not contain the prototype RBD, the neutralizing antibody titers against prototype strain elicited by “BBIBP-CorV+mos-tri-RBD” were no less than those elicited by “BBIBP-CorV+homo-tri-RBD”, both of which were significantly higher than those induced by“BBIBP-CorV+BBIBP-CorV” vaccinations (Figure 3B). Our results indicated that mos-tri-RBD was highly immunogenic as a booster dose against the prototype SARS-CoV-2 strain.

Compared with the prototype pseudo-virus, the neutralizing antibody GMTs against Omicron variant, including lineages BA.1, BA.2 and BA.3, elicited by “BBIBP-CorV+BBIBP-CorV” vaccination were significantly reduced (Figure 3B), suggesting high immune escape capability of Omicron variant from the immunity offered by BBIBP-CorV. In comparison to homologous booster of BBIBP-CorV, the “BBIBP-CorV+homo-tri-RBD” vaccination improved the neutralizing antibody GMTs against Omicron BA.1, BA.2 and BA.3 pseudo-viruses from 383, 152 and 81 to 932, 347 and 317, respectively, but these values were also significantly lower than the value against the prototype strain (Figure 3B). Furthermore, remarkably enhanced neutralizing antibody titers against Omicron were elicited by “BBIBP-CorV+mos-tri-RBD” vaccination, in which the neutralizing GMTs reached 3249, 1459 and 957 against BA.1, BA.2 and BA.3, respectively. The neutralizing GMTs boosted by mos-tri-RBD against the three lineages of Omicron variant were 3.5-fold, 4.2-fold and 3.0-fold higher than those boosted by homo-tri-RBD, and 8.5-fold, 9.6-fold and 11.8-fold higher than by BBIBP-CorV, respectively (Figure 3B). Mos-tri-RBD was immunogenically superior to homo-tri-RBD and BBIBP-CorV as a booster vaccine for BBIBP-CorV recipients against Omicron variant.

Considering that mos-tri-RBD also contains the key mutations from other SARS-CoV-2 variants with potential immune evasion ability, we then evaluate whether the mos-tri-RBD booster induced higher neutralizing responses than homo-tri-RBD and BBIBP-CorV against other pseudo-typed SARS-CoV-2 VOCs and VOIs, including Alpha, Beta, Delta, Gamma, Lambda and Mu variants. Pseudo-virus neutralization assays showed that for most of the tested variants, the neutralizing antibody titers in the sera elicited by “BBIBP-CorV+mos-tri-RBD” vaccination were higher than those by “BBIBP-CorV+mos-tri-RBD” and “BBIBP-CorV+BBIBP-CorV”. Especially, for Beta, Gamma, Delta, Lambda and Mu variants, the neutralizing antibody GMTs were increased 8.1-fold, 5.1-fold, 9.1-fold, 9.1-fold and 5.6-fold, respectively, for “BBIBP-CorV+mos-tri-RBD” vaccination compared to “BBIBP-CorV+BBIBP-CorV” vaccination, and 4.5-fold, 2.6-fold, 2.4-fold, 2.4-fold and 3.2-fold, respectively, compared to “BBIBP-CorV+homo-tri-RBD” vaccination (Figure 3B). These results indicated that mos-tri-RBD as a booster dose significantly improved the immunogenicity against not only Omicron variant but also other potentially immune-evasive SARS-CoV-2 variants. Mos-tri-RBD may act as a booster vaccine with broad-neutralization activities.

### Mos-tri-RBD as a booster dose induced cross-neutralization against the live viruses of SARS-CoV-2 prototype, Omicron, Beta and Delta strains

The significantly higher neutralizing activities boosted by mos-tri-RBD against multiple SARS-CoV-2 variants, including Omicron (BA.1), Beta and Delta, were further verified by using live virus neutralization assays. As a comparison, the neutralizing response against prototype strain was also tested.

Regarding the prototype strain, live virus neutralization assays showed that booster vaccinations with the three vaccines all elicited strong neutralization activities compared to adjuvant control. In line with the results of pseudo-virus neutralization assay, the live-virus neutralizing antibody titers induced by “BBIBP-CorV+mos-tri-RBD” and “BBIBP-CorV+homo-tri-RBD” vaccinations were significantly greater than those by “BBIBP-CorV+BBIBP-CorV” (Figure 4). The results demonstrated that mos-tri-RBD and homo-tri-RBD were more immunogenic than BBIBP-CorV against the prototype SARA-CoV-2 strain.

**Figure 4.**
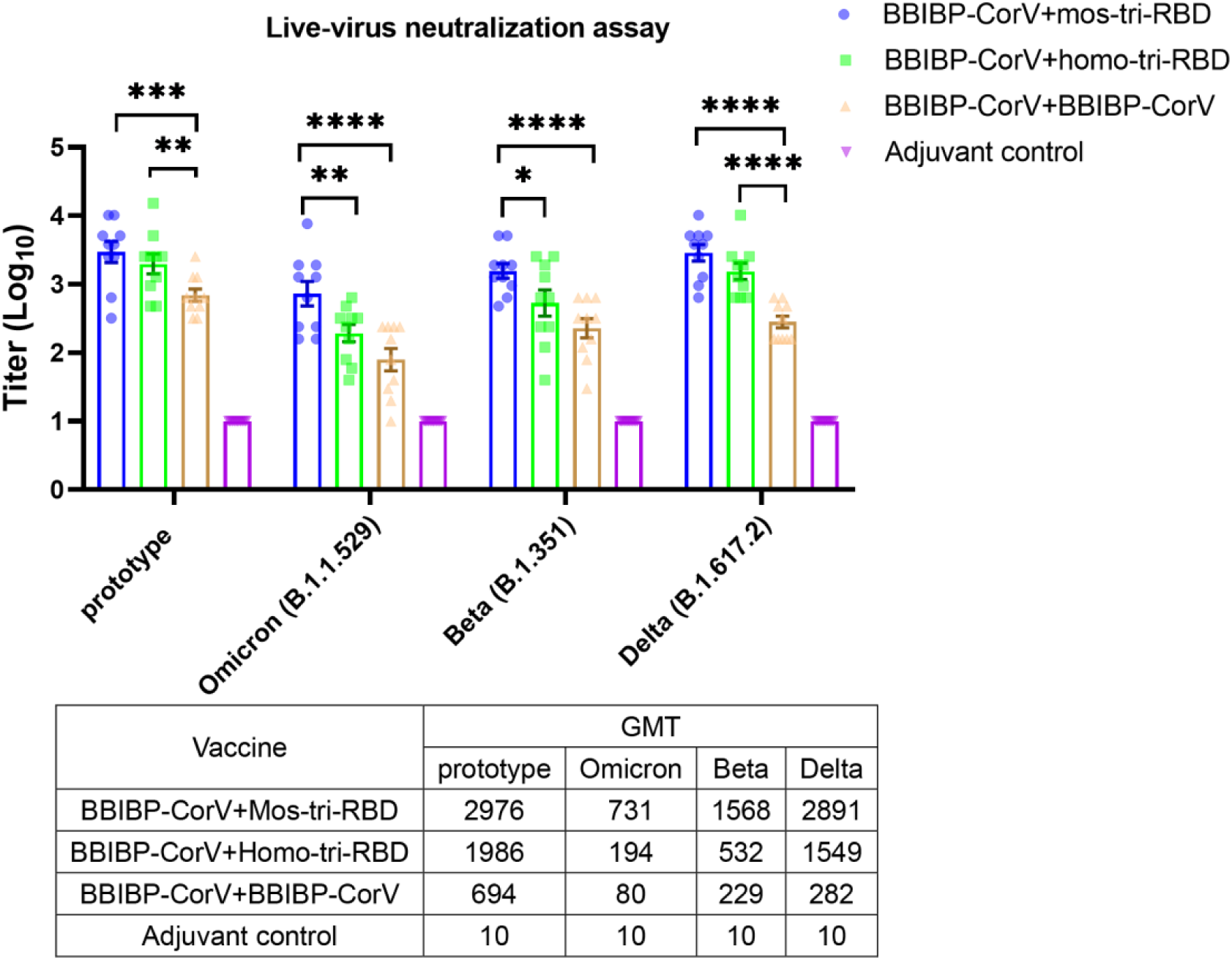
Evaluation of the cross-reactive immunogenicity of mos-tri-RBD as a booster shot against multiple SARS-CoV-2 strains, including prototype, Omicron, Beta and Delta strains, using live-virus neutralization assay. Neutralizing antibody titers elicited by “BBIBP-CorV+mos-tri-RBD” vaccination compared with those elicited by “BBIBP-CorV+homo-tri-RBD” and “BBIBP-CorV+BBIBP-Corv” vaccinations against the live-viruses of SARS-CoV-2 Omicron as well as other immune-evasive variants. Data are presented as mean ± SEM. One-way ANOVA followed by the LSD t-test was used for the comparison of data between different groups. *P<0.05, **P<0.01, ****P<0.0001. GMT values are displayed in the lower part of the figure. **Figure 4 – source data 1** **Individual data of live-virus neutralizing antibody titers against several SARS-CoV-2 circulating strains elicited by “BBIBP-CorV+mos-tri-RBD” vaccination compared with those elicited by “BBIBP-CorV+homo-tri-RBD” and “BBIBP-CorV+BBIBP-CorV” vaccinations**.

The neutralizing antibody response against Omicron variant boosted by BBIBP-CorV was dramatically reduced compared with that against prototype virus, suggesting significant immune evasion of Omicron from the homologous BBIBP-CorV booster vaccination. Compared with “BBIBP-CorV+BBIBP-CorV” vaccination, the neutralizing antibody GMT against Omicron variant in the “BBIBP-CorV+homo-tri-RBD” immunization group was increased from 80 in to 194, however, the value was also significantly lower than that against the prototype strain. Furthermore, similar to the results of pseudo-virus neutralization assay, “BBIBP-CorV+mos-tri-RBD” vaccination elicited remarkably improved live-virus neutralizing antibodies with a GMT value of 731 (Figure 4). Compared to booster vaccinations of BBIBP-CorV and homo-tri-RBD, boosting with mos-tri-RBD induced 9.1-fold and 3.8-fold higher Omicron-specific neutralizing antibodies, respectively, which provides an effective booster vaccine against Omicron variant. Similar results were also observed for Beta and Delta variants. In “BBIBP-CorV+BBIBP-CorV” vaccination groups, both Beta and Delta variants exhibited less sensitivity to neutralization by the sera from the immunized rats. While, the neutralizing antibody GMT against Beta variant elicited by “BBIBP-CorV+homo-tri-RBD” was increased from 229 to 532 (2.3-fold), and further improved to 1568 (6.8-fold) by “BBIBP-CorV+mos-tri-RBD” vaccination (Figure 4). For Delta variant, the neutralizing antibody GMT boosted by mos-tri-RBD and homo-tri-RBD was 5.5-fold and 10.3-fold, respectively, higher than that boosted by BBIBP-CorV (Figure 4). These results implied that mos-tri-RBD was immunologically superior to homo-tri-RBD and BBIBP-CorV as a booster dose against Omicron and other immune-evasive SARS-CoV-2 variants.

In summary, both pseudo- and live-virus neutralization assays demonstrated that the designed mos-tri-RBD could serve as an effective booster vaccine to elicit potent and cross-reactive immunity against not only prototype virus but also Omicron and other variants.

## Discussion

Facing the severe pandemic of SARS-CoV-2 variants, especially Omicron, WHO has recommended updating the composition of current COVID vaccines and developing multivalent or cross-protective vaccines against the variants (https://www.who.int/news/item/). Here, we designed a mosaic-type tri-RBD (mos-tri-RBD), which harbors the key mutations derived from Omicron and other immune-evasive variants, to broaden the immune response to SARS-CoV-2 variants. The immunogenicity of the designed mosaic-type vaccine candidate was assessed in rats, and live-virus neutralization assays showed that mos-tri-RBD elicited broad-spectrum neutralizing antibody response against multiple SARS-CoV-2 variants including Omicron. Considering that large-scale populations in the world have completed the primary series of vaccination, the immunogenicity of the designed mos-tri-RBD was also evaluated as a booster dose following the vaccination of BBIBP-CorV. Tests in rats showed that mos-tri-RBD booster vaccination elicited remarkably improved neutralizing antibodies against not only Omicron but also other SARS-CoV-2 VOCs and VOIs, which was immunogenically superior to the booster vaccinations of homo-tri-RBD and BBIBP-CorV. Thus, mos-tri-RBD may serve as a broad-neutralizing vaccine candidate, which could be used alone or as a booster shot in combating SARS-CoV-2 variants including Omicron.

A commonly used strategy for the development of broad-spectrum vaccines is to produce polyvalent vaccines that contain multiple strain-specific monovalent components. Considering the potential immune escape capability of Beta variant, several studies have designed the Beta-specific COVID-19 vaccines and applied combining with the anti-prototype ones to broaden immune response (Wu et al., 2021a; Callaway and Ledford, 2021; Logue et al., 2021). Targeting the Omicron strain with stronger immune evasion ability, several vaccine manufacturers have announced the update of the composition of their COVID-19 vaccines to provide effective protection against Omicron (Cohen, 2021). However, this conventional strategy for multivalent vaccine development is time-consuming and cost-expensive. In the present work, we provided a more promising strategy for the development of broad-spectrum vaccines against SARS-CoV-2, i.e., the construction of mosaic-type vaccines which incorporate multiple antigens and key mutations derived from different variants into a single hybrid immunogen. The mosaic strategy has been successfully applied to the development of broad-spectrum vaccines for HIV, coronaviruses and influenza (Barouch et al., 2010; Cohen et al., 2021; Kanekiyo et al., 2019), and several studies have demonstrated that mosaic-type immunogen elicited superior B cell responses both in quantity and quality compared to the homotypic immunogens (Kanekiyo et al., 2019). Our results also showed that the constructed mos-tri-RBD not only strengthens but also broadens neutralizing response against SARS-CoV-2.

In the designed mos-tri-RBD, except for the mutations from Omicron variant, the other integrated mutations were all in the top 10 most observed RBD mutations (https://weilab.math.msu.edu/MutationAnalyzer/), indicating that these mutations not only largely contribute to immune evasion but also have a high chance of recurrence in future variants. Therefore, the potent and broad neutralizing response induced by mos-tri-RBD suggested that the designed vaccine candidate may also provide neutralization activities against the emerging SARS-CoV-2 variants in the future.

The construction of mosaic-type immunogen, which combines the key mutations relevant to immune evasion into a single molecule, provides an effective strategy to achieve broad-spectrum neutralization in a single-component vaccine. The mosaic strategy may also be used in the developments of mRNA- and DNA-based COVID-19 vaccines.

## Materials and Methods

### Key resources table

| Key Resources Table |  |  |  |  |
| --- | --- | --- | --- | --- |
| Reagent type (species) or resource | Designation | Source or reference | Identifiers | Additional information |
| strain, strain background (*SARS-CoV-2 virus*) | Live SARS-CoV-2 prototype virus (QD-01 strain) | National Institute for Viral Disease Control and Prevention, China CDC |  |  |
| strain, strain background (*SARS-CoV-2 virus*) | Live SARS-CoV-2 Omicron virus (NPRC 2.192100005 strain) | National Institute for Viral Disease Control and Prevention, China CDC |  |  |
| strain, strain background (*SARS-CoV-2 virus*) | Live SARS-CoV-2 Beta virus (GD84 strain) | National Institute for Viral Disease Control and Prevention, China CDC |  |  |
| strain, strain background (*SARS-CoV-2 virus*) | Live SARS-CoV-2 Delta virus (GD96 strain) | National Institute for Viral Disease Control and Prevention, China CDC |  |  |
| strain, strain background (*SARS-CoV-2 pseudo-virus*) | SARS-CoV-2 prototype, Omicron (BA.1), Omicron (BA.2), Omicron (BA.3), Alpha, Beta, Delta, Gamma, Lambda and Mu pseudo-viruses | Wang et al., 2022; Li et al., 2021; Zhang et al., 2021; Nie et al., 2020 |  |  |
| cell line (*CHO*) | CHO-K1 cell line | ATCC | Cat#CCL-61, RRID:CVCL_0214 |  |
| cell line (*Homo-sapiens*) | Huh-7 cells | JCRB | Cat#JCRB0403; RRID: CVCL_0336 |  |
| cell line (*Chlorocebus sabaeus*) | Vero cells | National Institute for Food and Drug Control (NIFDC), Beijing, China |  |  |
| biological sample (*Wistar rats*) | Serum samples from immunized Wistar rats | This paper |  | Freshly isolated from immunized rats |
| antibody | SARS-CoV-2 Spike Antibody, Omicron Reactive, mouse mAb MM117 | Sino Biological Inc., China | Cat#40592-MM117 |  |
| antibody | Goat anti-mouse IgG-HRP | ZSGB-BIO | Cat#ZB-2305; RRID: AB_2747415 |  |
| recombinant DNA reagent | Plasmid-SARS-CoV-2-mos-tri-RBD | This paper |  |  |
| peptide, recombinant protein | Recombinant mos-tri-RBD protein (mammalian cell-expressed) | This paper |  |  |
| peptide, recombinant protein | Recombinant monomeric his-tagged RBD of the prototype SARS-CoV-2 strain (Baculovirus-insect cell-expressed) | Sino Biological Inc., China | Cat#40592-V08B |  |
| peptide, recombinant protein | Recombinant monomeric his-tagged RBD of the Omicron (B.1.1.529) SARS-CoV-2 strain (HEK 293 cell-expressed) | Sino Biological Inc., China | Cat#40592-V08H121 |  |
| peptide,<br>recombinant<br>protein | Recombinant<br>hACE2<br>protein<br>(mammalian<br>cell-expressed) | Sino Biological<br>Inc., China | Cat#10108-H08H |  |
| commercial<br>assay or kit | SPR | BIACore 8k,<br>GE Healthcare | N/A |  |
| chemical<br>compound, drug | Aluminum<br>hydroxide<br>adjuvant | This paper | N/A |  |
| software,<br>algorithm | UCSF<br>Chimera | Chimera team<br>at University of<br>California | https://www.cgl.ucsf.edu/chimera/ |  |
| software,<br>algorithm | BIACore<br>Insight<br>Evaluation<br>Software | GE Healthcare |  |  |
| software,<br>algorithm | GraphPad<br>Prism<br>version 8 | GraphPad<br>Software | https://www.graphpad.com/ |  |

### Protein expression and purification

The mosaic-type trimeric RBD (mos-tri-RBD) protein was constructed through connecting three heterologous RBDs (amino acid 319-537 in S proteins) into a single chain, which co-assembled into a trimeric structure. The designed protein was transiently expressed in CHO cells and purified by chromatography combined with ultrafiltration, as described in our previous paper (Liang et al., 2022). Briefly, the sequence of the designed mos-tri-RBD was codon-optimized and synthesized. After adding signal peptide and Kozak sequences to N terminus, the construct was cloned into the PTT5 plasmid vector via the *Hin* dIII and *Not* I restriction sites. The generated plasmid was validated by gene sequencing and then transfected into the CHO cells. After culture of 10-12 days, the supernatant was collected. Then, protein sample was purified by using ion-exchange chromatography and hydrophobic chromatography, followed by ultrafiltration. During purification, the samples from the eluted peaks were analyzed by SDS-PAGE and size exclusion-high-performance liquid chromatography (SEC-HPLC). The homo-tri-RBD used in this study was stably expressed by CHO cells and purified following the similar processes described above.

### SPR assay

Surface plasmon resonance (SPR) assay was performed to quantify the binding avidity of the recombinant mos-tri-RBD to the receptor hACE2 using BIAcore 8K (GE Healthcare) with NTA chips. Firstly, the His-tagged hACE2 protein was immobilized onto the chip surface. Then, the purified mos-tri-RBD protein sample was serially diluted in HBS-T buffer (HBS buffer and 0.05% Tween20). The diluted samples were injected at a flow rate of 30 µL/min for 120 s, and then the HBS-T buffer was flowed over the chip surface to facilitate dissociation of the bound protein for an additional 120s. Subsequently, the sensor chip surface was regenerated by injecting 350 mM EDTA solution with a flow rate of 30 µL/min for 120s. The SPR signal response was monitored as a function of time. BIAcoreTM Insight Evaluation software was used to analyze the experimental data. The binding kinetics between mos-tri-RBD and hACE2 was calculated using a 1:1 binding model, and then the association and dissociation rate constants, i.e., *k*_a_ and *k*_d_, as well as the apparent dissociation constant *K*_D_ were obtained.

### Ethics statement

Animal experiments were approved by the Institutional Animal Care and Use Committee (IACUC) of the National Vaccine and Serum Institute (NVSI) and conducted under Chinese animal use guidelines.

### Rat immunization

To test the immunogenicity of mos-tri-RBD, 10 Wistar rats with half male and half female (purchased from Beijing Vital River Laboratory Animal Technology Co., Ltd., China) were immunized intramuscularly by two doses, three weeks apart, of mos-tri-RBD (10 μg per dose) mixed with 300μg aluminum hydroxide adjuvant. Another three groups of rats received two doses of homo-tri-RBD, BBIBP-CorV or adjuvant, respectively, with the same immunization regimen were used for comparison. On day 7 after full vaccination, sera from the immunized rats were collected.

To evaluate the immune efficacy of mos-tri-RBD as a booster dose, a total of 30 Wistar rats, half male and half female, were intramuscularly primed with a dose (4μg/dose) of the inactivated vaccine BBIBP-CorV. After 3 weeks, 10 rats were boosted with 10 μg mos-tri-RBD mixed with 300μg aluminum hydroxide adjuvant. The other 20 rats were boosted with a dose of homo-tri-RBD (10 μg antigen mixed with 300μg aluminum hydroxide) or BBIBP-CorV (4μg/dose), for comparison. Another 10 rats were vaccinated with two doses of adjuvant with the same immunization interval as control. The sera of all the immunized rats were collected on day 7 post-boosting immunization.

### ELISA

To verify the functionality of the recombinant protein mos-tri-RBD, enzyme-linked immunosorbent assay (ELISA) was employed to measure its binding activity with an RBD-specific monoclonal neutralizing antibody MM117 (purchased from Sino Biological Inc., China, Cat#40592-MM117) and the binding activities of the his-tagged monomeric prototype and Omicron RBDs with MM117 were also evaluated as control. Protein samples were prepared with the starting concentration of 1 µg/mL in carbonate buffer, and subjected to three-fold serial dilutions. The diluted samples were then pipetted into the wells of the ELISA plates with 100 µL per well followed by incubation at 2-8 °C overnight. After removing the coating solution and washing the plate three times with PBS containing 0.05% Tween 20 (PBST), the remaining protein-binding sites were blocked by blocking buffer at 37°C for two hours and the plate was again washed three times with PBST. The neutralizing antibody MM117 was prepared with the working concentration of 1 µg/mL, which was added to the plate with 100 µL per well and incubated at 37°C for one hour. After washing three times with PBST, the plate was incubated with HRP-conjugated goat anti-mouse IgG antibody at 37°C for one hour. Then, color reaction was developed with 50 µL tetramethylbenzidine (TMB) and 50 µL hydrogen peroxide solutions. After 5 min, color reactions were stopped with sulfuric acidic solution, and the optical density (OD) was measured both at 450 nm and 630 nm using the microplate reader. The difference in OD values at 450 nm and 630 nm, i.e. OD_450/630nm_, was obtained to evaluate the specific binding activity of mos-tri-RBD with the neutralizing antibody MM117.

### Pseudo-virus neutralization assay

Neutralizing antibody levels in the sera from the immunized rats were detected by using pseudo-virus neutralization assay as described previously (Liang et al., 2022). In order to assess the cross-neutralization activities elicited by the vaccine candidate, we conducted neutralization assays against 10 pseudo-typed SARS-CoV-2 viruses, including prototype, Omicron (BA.1), Omicron (BA.2), Omicron (BA.3), Alpha, Beta, Delta, Gamma, Lambda and Mu strains. The pseudo-viruses were prepared by using the methods described in the previously published studies (Zhang et al., 2022; Li et al., 2021; Zhang et al., 2021; Nie et al., 2020). In short, the gene sequences of the S protein of SARS-CoV-2 variants were codon-optimized and synthesized, which were then cloned into pcDNA3.1 plasmid vector. The constructed plasmids encoding the S protein of SARS-CoV-2 variants were transfected into HEK293T cells, which were simultaneously infected with G*ΔG-VSV. After 24 hours, the culture supernatants were collected and filtered with 0.45μm membrane filters to obtain the VSV-based pseudo-typed SARS-CoV-2 variants. The TICD_50_ of the pseudo-viruses was determined by using HuH-7 cells.

To evaluate the pseudo-virus neutralization activities of the serum samples, the sera were firstly 1:40 diluted, followed by a 5-fold serial dilution with the cell culture medium. The pseudo-typed SARS-CoV-2 strains were also diluted to the titer of 1.3×10^4^ TCID_50_ per mL. Then, 50 µL diluted serum was mixed with an equal volume of pseudo-virus in the well of the plates, and incubated at 37 °C and 5% CO_2_ for 1 h. 50 µL culture medium mixed with 50 µL pseudo-virus was used as a control, and 100 µL per well medium without adding pseudo-virus served as a blank control. Subsequently, 100 µL trypsin-treated HuH-7 cells with the density of 2×10^5^ per mL were added into the well of the plates, and incubated at 37 °C and 5% CO_2_ for 20 ∼ 24 hours. After that, the cells were lysed and the luminescence signals were detected by microplate luminometer using luciferase substrate. The neutralizing antibody titer was determined as the reciprocal of the serum dilution causing 50% reduction in relative light units (RLUs), which was calculated using the Reed-Muench method. The titer for the serum below the limit of detection was set to half value of the detection limit.

### Live virus neutralization assay

A total of four SARS-CoV-2 live virus strains, including the prototype (QD-01 strain), Omicron (NPRC 2.192100005), Beta (GD84 strain) and Delta (GD96 strain), were tested in live virus neutralization assays, which was performed in the BSL3 facility of National Institute for Viral Disease Control and Prevention, Chinese Center for Disease Control and Prevention (China CDC), Beijing, China. Serum samples were heat-inactivated and serially diluted, which were then mixed with an equal volume of SARS-CoV-2 solution containing 100 TCID_50_ of live virus. The serum-virus mixed solution was incubated at 37 °C for 2 h. After incubation, Vero cell suspension with a density of 2×10^5^ per mL was added into the mixture. Cell-only and virus-only wells were also included in the plate as controls. The plates were incubated at 37 °C for 5 to 7 days. Then, inhibition of the serum on the infection of live virus was observed. Neutralizing antibody titer was reported as the reciprocal of the serum dilution for 50% protection against viral infection.

### Quantification and statistical analysis

One-way ANOVA with the LSD t-test method was used for the comparison of data from multiple groups, and Student’s t-test was used for statistical analysis between two groups. *P<0.05, **P<0.01, ***P<0.001, ****P<0.0001. Details can be found in the figure legend.

## Data availability

Figure 1 – Source Data 1, Figure 1 – Source Data 2, Figure 2 – Source Data 1, Figure 3 – Source Data 1, and Figure 4 – Source Data 1.

## Author contributions

Conceptualization: QML, JingZ, JGS, YL

Methodology: QML, GZW, JingZ, JGS, ZBH, YL

Investigation: ZBH, YL, KX, XFZ, YQJ, JWH, XL, YNH, LFD, JJW, XYL

Validation: SS, ZHL, ZML, NL, FJS, XZ

Resources: GZW, WJH, KX, HW, JinZ

Visualization: JGS, ZBH, YL

Supervision: QML, JingZ, GZW

Writing—original draft: JGS, ZBH, SS, YL, JingZ

Writing—review & editing: QML, GZW, WJH

## Acknowledgments

This work was supported by National Vaccine and Serum Institute (KTZC1900026C).

## References

Barouch DH, O’Brien KL, Simmons NL, King SL, Abbink P, Maxfield LF, Sun YH, Porte AL, Riggs AM, Lynch DM, Clark SL, Backus K, Perry JR, Seaman MS, Carville A, Mansfield KG, Szinger JJ, Fischer W, Muldoon M, Korber B (2010) Mosaic HIV-1 vaccines expand the breadth and depth of cellular immune responses in rhesus monkeys. Nature Medicine 16, 319–323.

Callaway E (2021) Omicron likely to weaken COVID vaccine protection. Nature 600:367–368.

Callaway E, Ledford H (2021) How to redesign COVID vaccines so they protect against variants. Nature 590:15–16.

Cao Y, Wang J, Jian F, Xiao T, Song W, Yisimayi A, Huang W, Li Q, Wang P, An R, Wang J, Wang Y, Niu X, Yang S, Liang H, Sun H, Li T, Yu Y, Cui Q, Liu S, Yang X, Du S, Zhang Z, Hao X, Shao F, Jin R, Wang X, Xiao J, Wang Y, Xie XS (2022) Omicron escapes the majority of existing SARS-CoV-2 neutralizing antibodies. Nature 602:657–663.

Cele S, Jackson L, Khoury DS, Khan K, Moyo-Gwete T, Tegally H, San JE, Cromer D, Scheepers C, Amoako DG, Karim F, Bernstein M, Lustig G, Archary D, Smith M, Ganga Y, Jule Z, Reedoy K, Hwa SH, Giandhari J, Blackburn JM, Gosnell BI, Karim SSA, Hanekom W, NGS-SA, COMMIT-KZN Team, von Gottberg A, Bhiman JN, Lessells RJ, Moosa MYS, Davenport MP, de Oliveira T, Moore PL, Sigal A (2022) Omicron extensively but incompletely escapes Pfizer BNT162b2 neutralization. Nature 602:654–656.

Cohen AA, Gnanapragasam PNP, Lee YE, Hoffman PR, Ou S, Kakutani LM, Keeffe JR, H. Wu Hj, Howarth M, West AP, Barnes CO, Nussenzweig MC, Bjorkman PJ (2021) Mosaic nanoparticles elicit cross-reactive immune responses to zoonotic coronaviruses in mice. Science 371:735–741.

Cohen J (2021) Omicron sparks a vaccine strategy debate. Science 374:1544–1545.

Dejnirattisai W, Zhou D, Supasa P, Liu C, Mentzer AJ, Ginn HM, Zhao Y, Duyvesteyn HME, Tuekprakhon A, Nutalai R, Wang B, López-Camacho C, Slon-Campos J, Walter TS, Skelly D, Clemens SAC, Naveca FG, Nascimento V, Nascixmento F, da Costa CF, Resende PC, Pauvolid-Correa A, Siqueira MM, Dold C, Levin R, Dong T, Pollard AJ, Knight JC, Crook D, Lambe T, Clutterbuck E, Bibi S, Flaxman A, Bittaye M, Belij-Rammerstorfer S, Gilbert SC, Carroll MW, Klenerman P, Barnes E, Dunachie SJ, Paterson NG, Williams MA, Hall DR, Hulswit RJG, Bowden TA, Fry EE, Mongkolsapaya J, Ren J, Stuart DI, Screaton GR (2021) Antibody evasion by the P.1 strain of SARS-CoV-2. Cell 184:2939-2954.e9.

Edara VV, Norwood C, Floyd K, Lai L, Davis-Gardner ME, Hudson WH, Mantus G, Nyhoff LE, Adelman MW, Fineman R, Patel S, Byram R, Gomes DN, Michael G, Abdullahi H, Beydoun N, Panganiban B, McNair N, Hellmeister K, Pitts J, Winters J, Kleinhenz J, Usher J, O’Keefe JB, Piantadosi A, Waggoner JJ, Babiker A, Stephens DS, Anderson EJ, Edupuganti S, Rouphael N, Ahmed R, Wrammert J, Suthar MS (2021) Infection-and vaccine-induced antibody binding and neutralization of the B.1.351 SARS-CoV-2 variant. Cell Host & Microbe 29:516-521.e3.

Kanekiyo M, Joyce MG, Gillespie RA, Gallagher JR, Andrews SF, Yassine HM, Wheatley AK, Fisher BE, Ambrozak DR, Creanga A, Leung K, Yang ES, Boyoglu-Barnum S, Georgiev IS, Tsybovsky Y, Prabhakaran MS, Andersen H, Kong WP, Baxa U, Zephir KL, Ledgerwood JE, Koup RA, Kwong PD, Harris AK, McDermott AB, Mascola JR, Graham BS (2019) Mosaic nanoparticle display of diverse influenza virus hemagglutinins elicits broad B cell responses. Nature Immunology 20:362–372.

Kupferschmidt K, Vogel G (2021) How bad is Omicron? Some clues are emerging. Science 374:1304–1305.

Lazarevic I, Pravica V, Miljanovic D, Cupic M (2021) Immune evasion of SARS-CoV-2 emerging variants: what have we learnt so far? Viruses 13:1192.

Li Q, Nie J, Wu J, Zhang L, Ding R, Wang H, Zhang Y, Li T, Liu S, Zhang M, Zhao C, Liu H, Nie L, Qin H, Wang M, Lu Q, Li X, Liu J, Liang H, Shi Y, Shen Y, Xie L, Zhang L, Qu X, Xu W, Huang W, Wang Y (2021) SARS-CoV-2 501Y.V2 variants lack higher infectivity but do have immune escape. Cell 184:2362–2371.

Liang Y, Zhang J, Yuan RY, Wang MY, He P, Su JG, Han ZB, Jin YQ, Hou JW, Zhang H, Zhang XF, Shao S, Hou YN, Liu ZM, D. LF, Shen FJ, Zhou WM, Xu K, Gao RQ, Tang F, Lei ZH, Liu S, Zhen W, Wu JJ, Zheng X, Liu N, Chen S, Ma ZJ, Zheng F, Ren SY, Hu ZY, Huang WJ, Wu GZ, Ke CW, Li QM (2022) Design of a mutation-integrated trimeric RBD with broad protection against SARS-CoV-2. Cell Disc. 8:17.

Lipsitch M, Krammer F, Regev-Yochay G, Lustig Y, Balicer RD (2022) SARS-CoV-2 breakthrough infections in vaccinated individuals: measurement, causes and impact. Nature Reviews Immunology 22:57–65.

Liu H, Wei P, Zhang Q, Aviszus K, Linderberger J, Yang J, Liu J, Chen Z, Waheed H, Reynoso L, Downey GP, Frankel SK, Kappler J, Marrack P, Zhang G (2021) The Lambda variant of SARS-CoV-2 has a better chance than the Delta variant to escape vaccines. bioRxiv doi: 10.1101/2021.08.25.457692.

Liu L, Iketani S, Guo Y, Chan JFW, Wang M, Liu L, Luo Y, Chu H, Huang Y, Nair MS, Yu J, Chik KKH, Yuen TTT, Yoon C, To KKW, Chen H, Yin MT, Sobieszczyk ME, Huang Y, Wang HH, Sheng Z, Yuen KY, Ho DD (2022) Striking antibody evasion manifested by the Omicron variant of SARS-CoV-2. Nature 602:676–681.

Logue J, Johnson R, Patel N, Zhou B, Maciejewski S, Zhou H, Portnoff A, Tian JH, McGrath M, Haupt R, Weston S, Hammond H, Guebre-Xabier M, Dillen C, Plested J, Cloney-Clark S, Greene AM, Massare M, Glenn G, Smith G, Frieman M (2021) Immunogenicity and in vivo protection of a variant nanoparticle vaccine that confers broad protection against emerging SARS-CoV-2 variants. bioRxiv doi: 10.1101/2021.06.08.447631.

Lou F, Li M, Pang Z, Jiang L, Guan L, Tian L, Hu J, Fan J, FanH (2021) Understanding the secret of SARS-CoV-2 variants of concern/interest and immune escape. Frontiers in Immunology 12:744242.

Muik A, Wallisch AK, Sänger B, Swanson KA, Mühl J, Chen W, Cai H, Maurus D, Sarkar R, Türeci Ö, Dormitzer PR, Şahin U (2021) Neutralization of SARS-CoV-2 lineage B.1.1.7 pseudovirus by BNT162b2 vaccine-elicited human sera. Science 371:1152–1153.

Nie J, Li Q, Wu J, Zhao C, Hao H, Liu H, Zhang L, Nie L, Qin H, Wang M, Lu Q, Li X, Sun Q, Liu J, Fan C, Huang W, Xu M, Wang Y (2020) Quantification of SARS-CoV-2 neutralizing antibody by a pseudotyped virus-based assay. Nature Protools 15:3699–3715.

Pegu A, O’Connell SE, Schmidt SD, O’Dell S, Talana CA, Lai L, Albert J, Anderson E, Bennett H, Corbett KS, Flach B, Jackson L, Leav B, Ledgerwood JE, Luke CJ, Makowski M, Nason MC, Roberts PC, Roederer M, Rebolledo PA, Rostad CA, Rouphael NG, Shi W, Wang L, Widge AT, Yang ES, mRNA-1273 Study Group, Beigel JH, Graham BS, Mascola JR, Suthar MS, McDermott AB, Doria-Rose NA (2021) Durability of mRNA-1273 vaccine-induced antibodies against SARS-CoV-2 variants. Science 373:1372–1377.

Pettersen EF, Goddard TD, Huang CC, Couch GS, Greenblatt DM, Meng EC, Ferrin TE (2004) UCSF Chimera – a visualization system for exploratory research and analysis. J. Comput. Chem. 25:1605–1612.

Planas D, Bruel T, Grzelak L, Guivel-Benhassine F, Staropoli I, Porrot F, Planchais C, Buchrieser J, Rajah MM, Bishop E, Albert M, Donati F, Prot M, Behillil S, Enouf V, Maquart M, Smati-Lafarge M, Varon E, Schortgen F, Yahyaoui L, Gonzalez M, Sèze JD, Péré H, Veyer D, Sève A, Simon-Lorière E, Fafi-Kremer S, Stefic K, Mouquet H, Hocqueloux L, van der Werf S, Prazuck T, Schwartz O (2021) Sensitivity of infectious SARS-CoV-2 B.1.1.7 and B.1.351 variants to neutralizing antibodies. Nature Medicine 27:917–924.

Rössler A, Riepler L, Bante D, von Laer D, Kimpel J (2021) SARS-CoV-2 B.1.1.529 variant (Omicron) evades neutralization by sera from vaccinated and convalescent individuals. medRxiv doi:10.1101/2021.12.08.21267491.

Shen X, Tang H, Pajon R, Smith G, Glenn GM, Shi W, Korber B, Montefiori DC (2021) Neutralization of SARS-CoV-2 variants B.1.429 and B.1.351. The New England Journal of Medicine 384:2352–2354.

Torjesen I (2021) Covid-19: Omicron may be more transmissible than other variants and partly resistant to existing vaccines, scientists fear. BMJ 375:n2943.

Vaughan A (2021) Omicron emerges. New Scientist 252:7.

Wang P, Casner RG, Nair MS, Wang M, Yu J, Cerutti G, Liu L, Kwong PD, Huang Y, Shapiro L, Ho DD (2021) Increased resistance of SARS-CoV-2 variant P.1 to antibody neutralization. Cell Host & Microbe 29, 747-751.e4.

Widge AT, Rouphael NG, Jackson LA, Anderson EJ, Roberts PC, Makhene M, Chappell JD, Denison MR, Stevens LJ, Pruijssers AJ, McDermott AB, Flach B, Lin BC, Doria-Rose NA, O’Dell S, Schmidt SD, Neuzil KM, Bennett H, Leav B, Makowski M, Albert J, Cross K, Edara VV, Floyd K, Suthar MS, Buchanan W, Luke CJ, Ledgerwood JE, Mascola JR, Graham BS, Beigel JH, mRNA-1273 Study Group (2021) Durability of responses after SARS-CoV-2 mRNA-1273 vaccination. The New England Journal of Medicine 384:80–82.

Wu K, Choi A, Koch M, Elbashir S, Ma L, Lee D, Woods A, Henry C, Palandjian C, Hill A, Jani H, Quinones J, Nunna N, O’Connell S, McDermott AB, Falcone S, Narayanan E, Colpitts T, Bennett H, Corbett KS, Seder R, Graham BS, Stewart-Jones GBE, Carfi A, Edwards DK (2021a) Variant SARS-CoV-2 mRNA vaccines confer broad neutralization as primary or booster series in mice. Vaccine 39:7394–7400.

Wu K, Werner AP, Koch M, Choi A, Narayanan E, Stewart-Jones GBE, Colpitts T, Bennett H, Boyoglu-Barnum S, Shi W, Moliva JI, Sullivan NJ, Graham BS, Carfi A, Corbett KS, Seder RA, Edwards DK (2021b) Serum neutralizing activity elicited by mRNA-1273 vaccine. The New England Journal of Medicine 384:1468–1470.

Zhang L, Li Q, Liang Z, Li T, Liu S, Cui Q, Nie J, Wu Q, Qu X, Huang W, Wang Y (2022) The significant immune escape of pseudotyped SARS-CoV-2 variant Omicron. Emerging Microbes & Infections 11:1–5.

Zhang L, Cui Z, Li Q, Wang B, Yu Y, Wu J, Nie J, Ding R, Wang H, Zhang Y, Liu S, Chen Z, He Y, Su X, Xu W, Huang W, Wang Y (2021) Ten emerging SARS-CoV-2 spike variants exhibit variable infectivity, animal tropism, and antibody neutralization. Communications Biology. 4:1196.

